# Linear infrastructure and associated wildlife accidents create an ecological trap for an apex predator and scavenger

**DOI:** 10.1101/2024.04.20.590377

**Authors:** Navinder J Singh, Michelle Etienne, Göran Spong, Frauke Ecke, Birger Hörnfeldt

**Affiliations:** Department of Wildlife, Fish and Environmental Studies, Faculty of Forest Sciences, Swedish University of Agricultural Sciences, Umeå, 90183

**Keywords:** HIREC, animal ecology, animal movement, animal behaviour, integrated step selection function, wildlife traffic accidents, maladaptive behaviour, fitness, habitat selection

## Abstract

Animals can be caught in an “ecological trap” when they select for seemingly attractive habitats at the expense of their fitness. Such maladaptive behaviour is often a consequence of human induced rapid changes in animals’ natal environment such as building of energy and transportation infrastructure. We tested the ecological trap hypotheses for human created linear infrastructure on a widely distributed apex predator and a scavenger – the Golden eagle (*Aquila chrysaetos*), whose range spans across the entire northern hemisphere. Roads and railways create novel and attractive feeding subsidies through traffic induced mortality of other species, while powerline areas provide perching or nesting sites and scavenging opportunities from electrocuted or collision-killed birds. These conditions lead to negative demographic consequences for eagles. We used integrated step selection functions for habitat selection and movement behaviour with ten years of data from 74 GPS-tracked Golden Eagles (36 adult and 38 immature) in Fennoscandia. To measure habitat attractiveness, we use wildlife traffic accident statistics on major wildlife species including the eagles, and mortality of five GPS-tracked eagles to show demographic consequences. Eagles selected for linear features all year round and across entire study region. Individuals also searched and sat alongside roads and railway lines more frequently. Immature eagles selected roads and railway sites more consistently compared to adults and showed learning behaviour with age. We discuss implications of these findings for conservation and population ecology of apex predators and scavengers and their potential evolutionary implications. We suggest that rapid removal of carcasses from roads and tracks is urgently needed to avoid this trap for many raptor and scavenger species throughout the world and develop methods and approaches to reduce wildlife traffic accidents all together.

## Introduction

Animals rely on a variety of cues (e.g. certain habitat features, sound, light, smell) for selecting habitats, navigation, foraging and mate choice with the aim to increase their survival and reproduction and eventually, their fitness (Darwin, 1859). Human modifications of the natural ecosystems are rapidly affecting, and at times, destroying these cues that animals use (Sih, 2013). As a result, animals may select habitats which may seem attractive, but result in lower fitness, thus falling into an ‘ecological trap’ (Gates & Gysel, 1978; Robertson et al., 2013). Ecological traps have been observed to have reduced fitness in several animal groups such as insects (Horváth et al., 2010), reptiles (Hawlena et al., 2010), fish (Jeffres & Moyle, 2012), birds (Remeš, 2003) and mammals (Lamb et al., 2017). Sea turtle hatchlings for instance are driven by reflected moonlight to move towards the ocean under natural circumstances but might be caught in an ecological trap when moving to a light polluted beach instead, thereby increasing mortality risk (Witherington, 1997).

For an ecological trap to exist, three criteria must be fulfilled: when comparing the trap habitat to surrounding habitats (i) the trap habitat is preferred (severe trap) or animals equally select both habitats (equal preference trap), (ii) individual fitness is lower in the trap habitat, (iii) animals actively move into the trap habitat (Hale et al., 2015; Robertson & Hutto, 2006). Hawlena et al. (2010) demonstrated an equal preference ecological trap where individuals of the lizard *Acanthodactylus beershebensis* equally selected for natural and a human modified habitat, in which predator exposure was higher, resulting in increased mortality. Similarly, a population of blackcaps (*Sylvia atricapilla*) selected a human-modified landscape with newly introduced plant species over their natural breeding habitat, resulting in a severe ecological trap with lower breeding success (Remeš, 2003). A widespread human modification of ecosystem with high potential of forming ecological trap is the establishment of linear infrastructure comprising roads, railways and powerlines. This creates an attractive, open and predictable hunting and scavenging habitat (through carrion from traffic accidents and electrocution of forage species from powerlines) but with risks from collisions, barriers and electrocution on the consumers (Harris & Scheck, 1991; Seiler, 2001).

Large predators and scavengers such as eagles and vultures are especially vulnerable to potential ecological traps created by linear infrastructure, because they move effectively over landscapes, are long lived and have a strong ability to learn, lack natural predators, and thus can fail to perceive novel risks (Ripple et al., 2014). Learning in raptors is mainly acquired through parental care and social interactions or predictability of food from supplementary feeding by humans (Cushing, 1944). Moreover, animals often select habitats similar to where they were born (i.e. natal habitat, Davis & Stamps, 2004). Hence, if immature individuals learn to use trap habitat as scavenging opportunity or recognize it as natal habitat, they may consistently use this habitat over years, and have long term negative effects on survival and future reproduction (Fletcher et al., 2015). A large soaring, long-lived raptor of conversation concern which is potentially vulnerable to ecological trap is the Golden Eagle (*Aquila chrysaetos*). Golden Eagles are widely distributed across the Holarctic (Watson, 2010), and listed as near threatened in Sweden (ArtDatabanken, 2015), a species needing special habitat conservation measures in the EU Birds Directive - Annex 1,European Union, 2009). As apex predators, Golden Eagles are opportunistic and also strongly depend on naturally fluctuating abundances of mountain hares (*Lepus timidus*) and different grouse species in the boreal forest (Moss et al., 2012; Tjernberg, 1981). In highly seasonal environments with harsh winters, eagles may use scavenging opportunities especially when prey abundance is low.

A recent study from Eisaguirre et al., (2020) found that Golden Eagles in Alaska selected roads and railways during migration periods and spent more time there compared to other habitats. In Sweden, eagles exhibit partial migration patterns mainly linked with age as well as winter conditions (Moss et al., 2014), with southward winter migration in late September to October and returning north in late April to early May (Singh et al., 2017). The Fennoscandian population uses old growth forests as breeding habitat and clear cuts and other open land with high prey detectability for hunting, while also using extensive scavenging opportunities (Singh et al., 2017; Tjernberg, 1981; Watson, 2010). In 2019 there were about 65000 reported fatal wildlife accidents on roads and railways in Sweden (primarily large mammals and eagles) with numbers increasing from 45000 in 2010 upto about 72000 (Swedish National Road Administration database, SNRAD - Nationella Viltolycksrådet, See Appendix). According to the National Traffic Management Agency (Trafikverket), 619 Golden Eagles and white-tailed sea eagles (*Haliaeetus albicilla*) died in traffic collisions during 2010-2023 (Figure 1a & 1c, and Appendix). Main causes of deaths in Golden Eagles recovered by the Swedish National Museum of Natural History, Stockholm, between 2003 - 2011 are attributed to collisions with trains (35,6 %), electrocution and powerline collisions (17,8 %) and starvation and other trauma that includes physical injuries (11,9 %), respectively (Ecke et al., 2017).

**Figure 1a.**
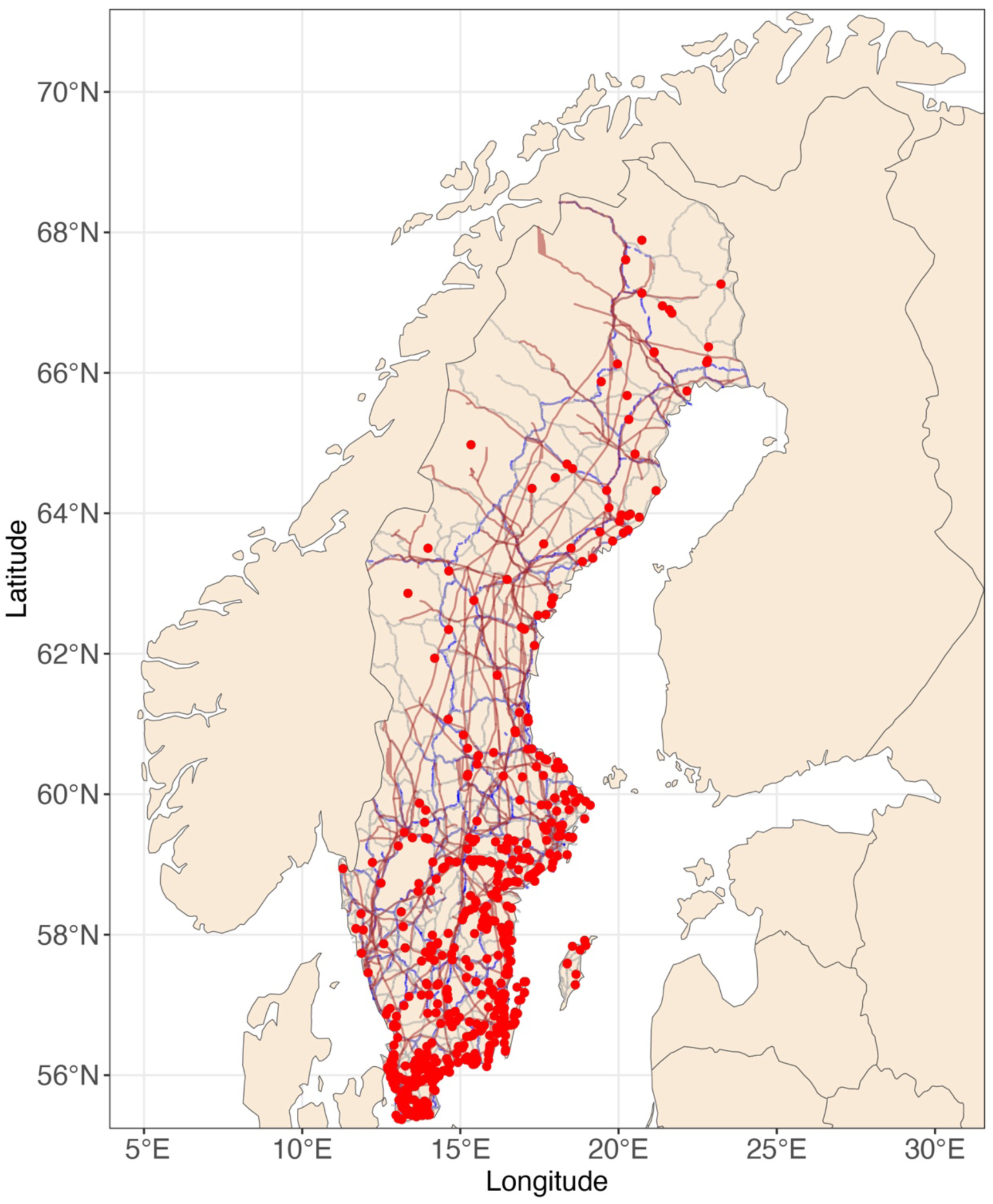
Locations of Eagle Traffic collisions (Red dots) recorded throughout Sweden during 2010-2023 (n=650) by the SNRAD. Only Primary Roads are shown, represented in grey, Railway lines in blue and the main Powerlines in brown.

**Figure 1b.**
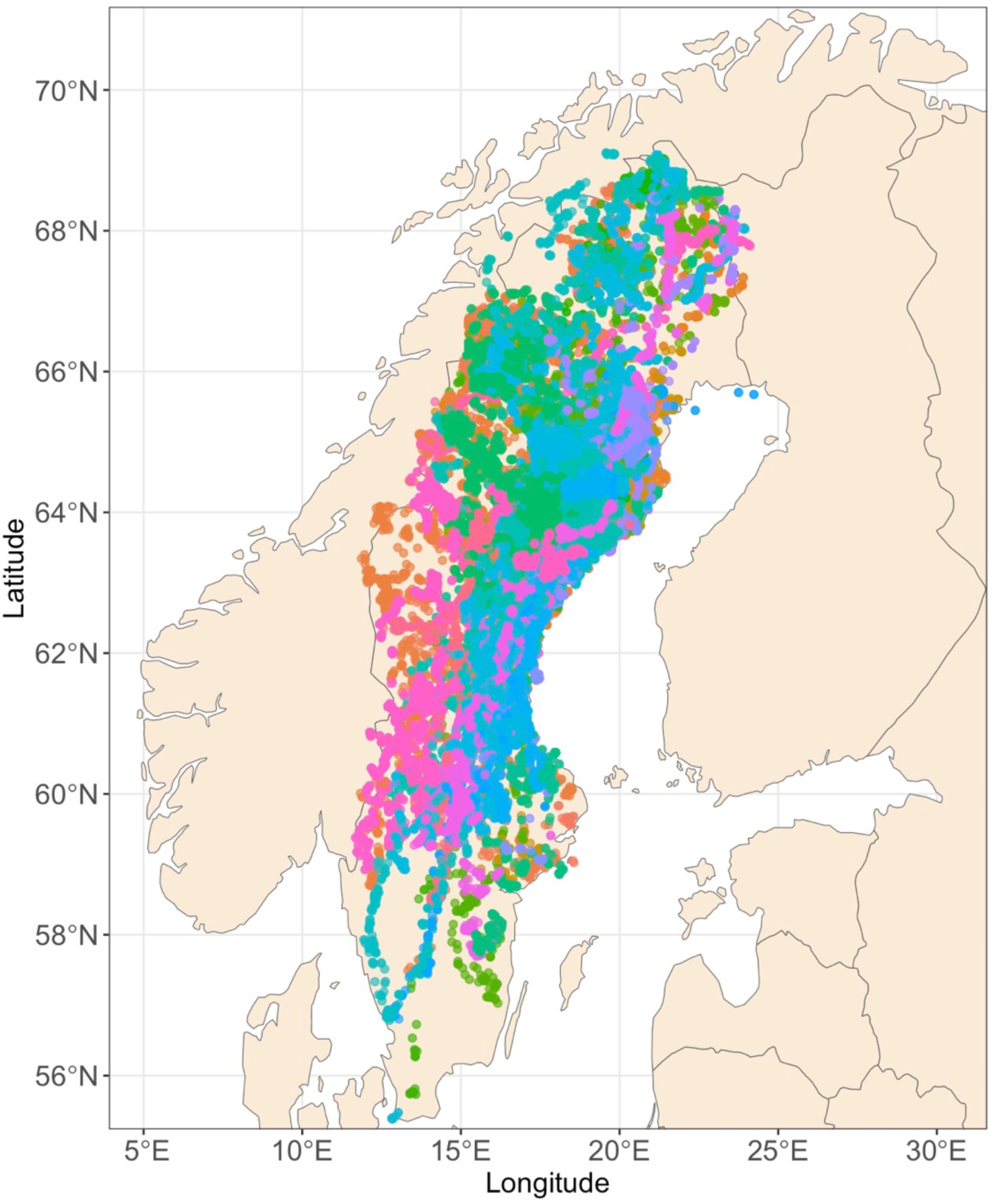
Multiannual movement of patterns of 74 studied Golden eagles within Sweden during 2010-2020 (n= 1031791). Individuals are represented by different coloured dots and only the positions within Sweden are shown.

**Figure 1c.**
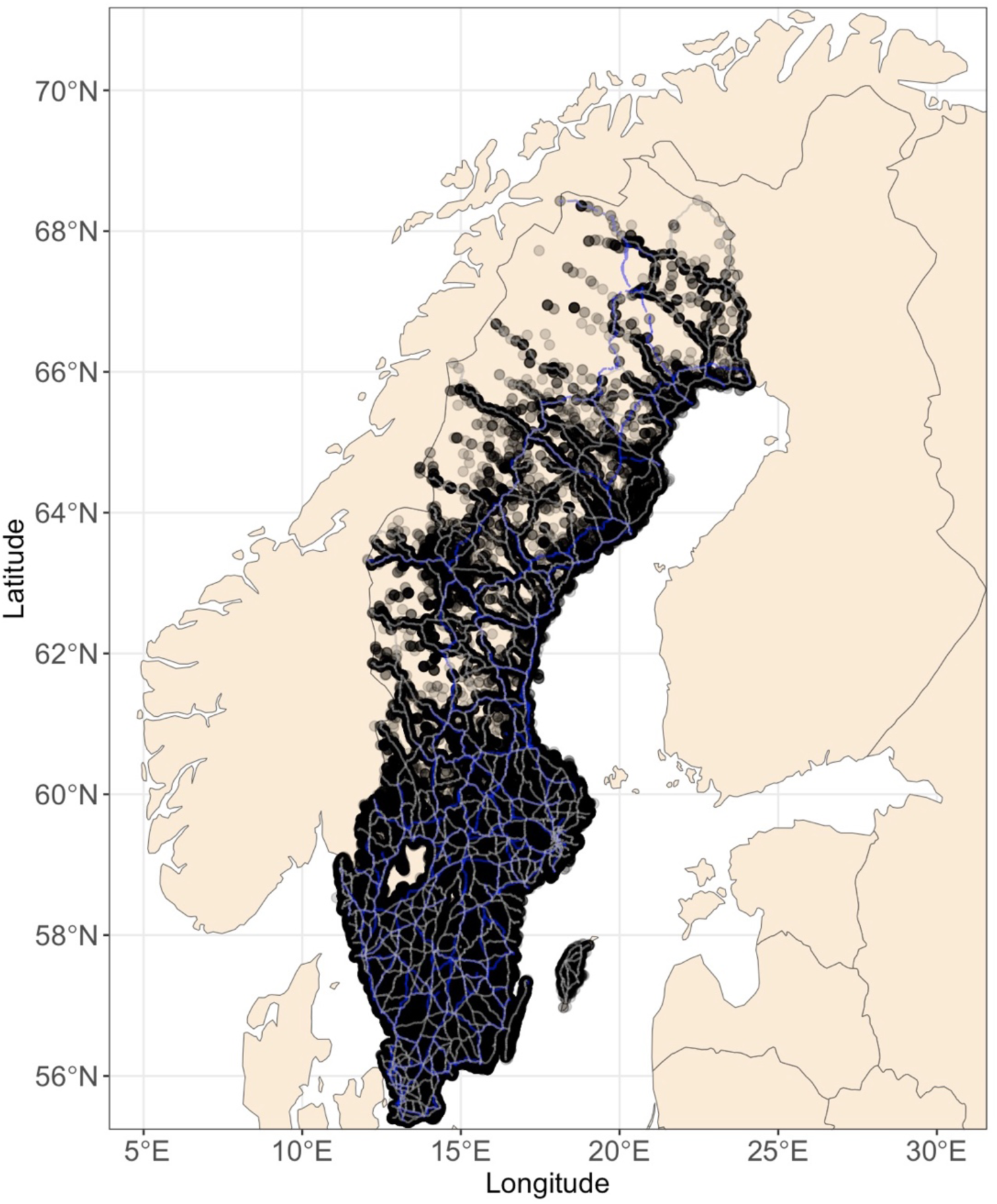
Locations of Wildlife Traffic Accidents (n=657249) recorded at SNRAD within Sweden during 2010-2023 overlaid on Roads and Railway lines, reflecting habitat attractiveness. Each dot represents a single instance. The species included are Moose, Roe deer, Red deer, Fallow deer, Wild boar, Bear, Wolf, Lynx, Wolverine and Otter.

Considering this large magnitude of mortality of eagles along linear infrastructure, in this study, we test if linear infrastructure creates an ecological trap for Golden Eagles. We made the following predictions based on the theory of ecological traps: (i) eagles consistently select for linear infrastructure (roads, railways and powerlines) as these features provide attractive scavenging opportunities from wildlife traffic accidents and electrocuted or collision-killed birds, while powerline poles provide perching sites due to good visibility for hunting in cleared powerline corridors. We predict that eagles have a high mortality in these habitats from traffic collisions and electrocution. Furthermore, we predict that (ii) eagles actively exhibit search behaviour for scavenging opportunities and have disproportionately higher frequency of sitting along the linear infrastructure; (iii) immature eagles select these habitats more than experienced adults due to their lower hunting success and finally, due to predictability of food abundance, (iv) over time immature eagles learn to use these habitats i.e. the strength of selection of these habitats increases with age.

## Methods and Study area

The study uses data from 74 GPS-tagged Golden Eagles (36 adults, 38 immatures) from Sweden within a study period of 11 years (2010 to 2020). Individual tracking periods ranged from one month to six years with a minimum of 500 relocations (Figure S1 and S2 Supplementary material). The study individuals ranged across most of Sweden (55 – 68°N, 12. – 23°E, Figure S1 Supplementary material). Adults were captured using remote controlled bownets (Bloom et al., 2007, 2015; Jackman et al., 1994) and tagged with solar-powered, backpack mounted global positioning systems (GPS) representing different transmitter types; in 2010-11: 75 g Microwave Telemetry Inc., USA and 140 g VectronicAerospace GmbH, Germany, and in 2014: 70 g Cellular Tracking Technologies, Inc., USA, with a maximum location error ranging from 10-18 m for all transmitters. Immatures were tagged as nestlings approximately two weeks prior to fledgling (Sandgren et al., 2014). All GPS tagging was conducted under Ethical Permits Nos. A 57-10, A 58-10, A 57-10A, A 33-13, and A11-2019 from the Swedish Agricultural Board - Jordbruksverket and Research Permit No. NV-07710-19 of The Swedish Environmental Protection Agency - Naturvårdsverket.

Forestry is the main land use across the Golden Eagle range in Sweden. The boreal forest landscape is characterised by cut-over and even-aged forests dominated by Norway Spruce (*Picea abies*) and Scots Pine (*Pinus sylvestris*) (Ecke et al., 2013; Esseen et al., 1997). The remaining landscape is a mix of forests interspersed with wetlands (lakes, streams and mires) and agricultural land (Helmfried, 1996). National distribution of roads, railways and powerlines and elevation and landcover data (100 m cell size) were based on raster layers in a geographic information system (GIS) provided by the Swedish mapping, cadastral and land registration authority (*Lantmäteriet*) and Swedish National Environmental Protection Agency (*Naturvårdsverket*). We refer to this linear infrastructure as ‘trap habitat’ throughout the study.

### Wildlife traffic accidents

We obtained monthly summaries of reported national wildlife traffic accidents at roads and railways for the years 2010-2023 from the Swedish National Road Administration database. White-tailed sea eagle and Golden eagle are summarized as one species because of difficulties to correctly identify the two eagle species after collision. Location data of eagles that died in traffic accidents were available from 2010-2023. Locations and timings of death were available for five out of six recovered GPS tagged Golden eagles from our project that died in wildlife traffic accidents.

## Data analyses

We applied an integrated step selection function (Avgar et al., 2016) using the package ‘amt’ for R (Signer et al., 2019) to investigate Golden Eagle habitat selection and behaviour around linear infrastructure. Locations were treated as linear steps between two consecutive relocations (Fortin et al., 2005; Thurfjell et al., 2014). Step selection functions test for habitat selection by conducting a conditional logistic regression comparing available to used habitat by including a random factor, while integrated step selection functions allow to include movement parameters into habitat selection analysis (Avgar et al., 2016), which reduce inferential bias (Forester et al., 2009) Eagle locations were resampled to an interval of 1 h ± 10 min to achieve a regular time interval. For each true eagle step (39747 in total) a set of random steps (n = 10) was created as a measure of availability. All step lengths (m) were assumed to follow a gamma distribution and all covariates were extracted at step end.

### Habitat Selection

The landcover data was classified into five categories, *viz*. ‘open land’, ‘forest’, ‘road and railway’, ‘water’ and ‘clear cut’ with reference to earlier studies on eagle biology (Singh et al., 2017). Open land was defined as ‘arable land’, ‘non-vegetated other open land’ and ‘vegetated other open land’ and used as intercept in the model, as no references for eagles strongly selecting or avoiding these types of land were found. Percentage of true eagle steps in open land were 10.3 %, 54.8 % in forest, 0.8 % at roads and railways, 0.6 % above water, 26.5 % in clear cuts and 6.8 % in other habitat like urban areas or artificial surfaces which were not included in habitat selection analysis. For further analyses, distance to linear infrastructure for each step was estimated as the Euclidean distance (m) to the nearest road, railway and powerline, respectively and tested in separate models due to correlation between types of infrastructure.

Seasons were defined as follows: Spring from March to May, Summer between June and August, Autumn between September and November and Winter between December and February. Percentage of true eagle steps in Spring were 33.2 %, 45.9 % in Summer, 17.4 % in Autumn and 3.6 % in Winter. Summer was used as the reference season to which the selection during other seasons was compared in the models.

### Behaviour

Eagle flight heights (m above ground level) for each position were calculated as the difference between the altitude reading obtained from GPS (m) and ground elevation data (m above sea level). Positions below 30 m were assumed as ‘sitting’ and above 30 m as ‘flying’ based on maximal boreal forest height (Larsen, 2013). Search and travel behaviour were estimated using the following method. We used the ‘amt’ package to estimate step lengths and turning angles. The cosine of the von Mises distributed turning angles reaches from −1 to 1 and can be used to describe movement direction, where positive values represent moving forwards from the previous location, zero represents a random walk and negative values represent moving backwards (Benhamou, 2006). We defined behaviour as ‘search’ when cosine of the turning angle was < 0 and step length < 1000 m. When searching for food, eagles are soaring (backwards movement) in small steps, while travel movements are forward with larger steps. Only flying positions (n = 182 044) were included in search behaviour analysis, and movements other than ‘search’ were treated as ‘travel’ to ensure model simplicity. The log of the step length was included in the models that tested behaviour and type of position as a modifier of the shape parameter of the underlying gamma distribution. As step length distribution differs between different behavioural modes, the log of the step length can be used to improve model efficiency (Avgar et al., 2016).

### Age effects and temporal pattens

For testing the effect of age on habitat selection, the results were compared between adults and immatures, with adults used as reference in the model. To investigate the learning behaviour, i.e. increase or stability of selection strength of trap habitat over time, individual eagle data were divided into eagle years, where the first year was defined as the next 365 days after tagging. All spatial and statistical analysis were performed in R (R version 3.6.1, R Core Team, 2019) and part of spatial data analysis in QGIS (QGIS version 3.42, QGIS Development Team, 2009).

## Results

### Selection of trap habitat

At the population level, eagles selected for roads and railways, clear cuts and forest indicated by a positive coefficient and a strong avoidance of water compared to open land indicated by a negative coefficient (Coefficient ± Standard Error, β_road+railway_ = 0.13 ± 0.06, β_clearcut_ = 0.99 ± 0.02, β_forest_ = 0.67 ± 0.02, β_water_ = - 2.24 ± 0.07, Table 1, Figure 2). Individuals showed a consistent temporal selection for roads, railways and powerlines throughout autumn, winter and spring compared to summer (β_road:autumn_ = - 0.38 ± 0.05, β_road:spring_ = - 0.38 ± 0.04, β_road:winter_ = - 0.42 ± 0.11, β_railway:autumn_ = - 0.28 ± 0.03, β_railway:spring_ = - 0.29 ± 0.03, β_railway:winter_ = - 1.162 ± 0.13, β_powerline:autumn_ = - 0.36 ± 0.05, β_powerline:spring_ = - 0.25 ± 0.03, β_powerline:winter_ = - 0.53 ± 0.12).

**Figure 2.**
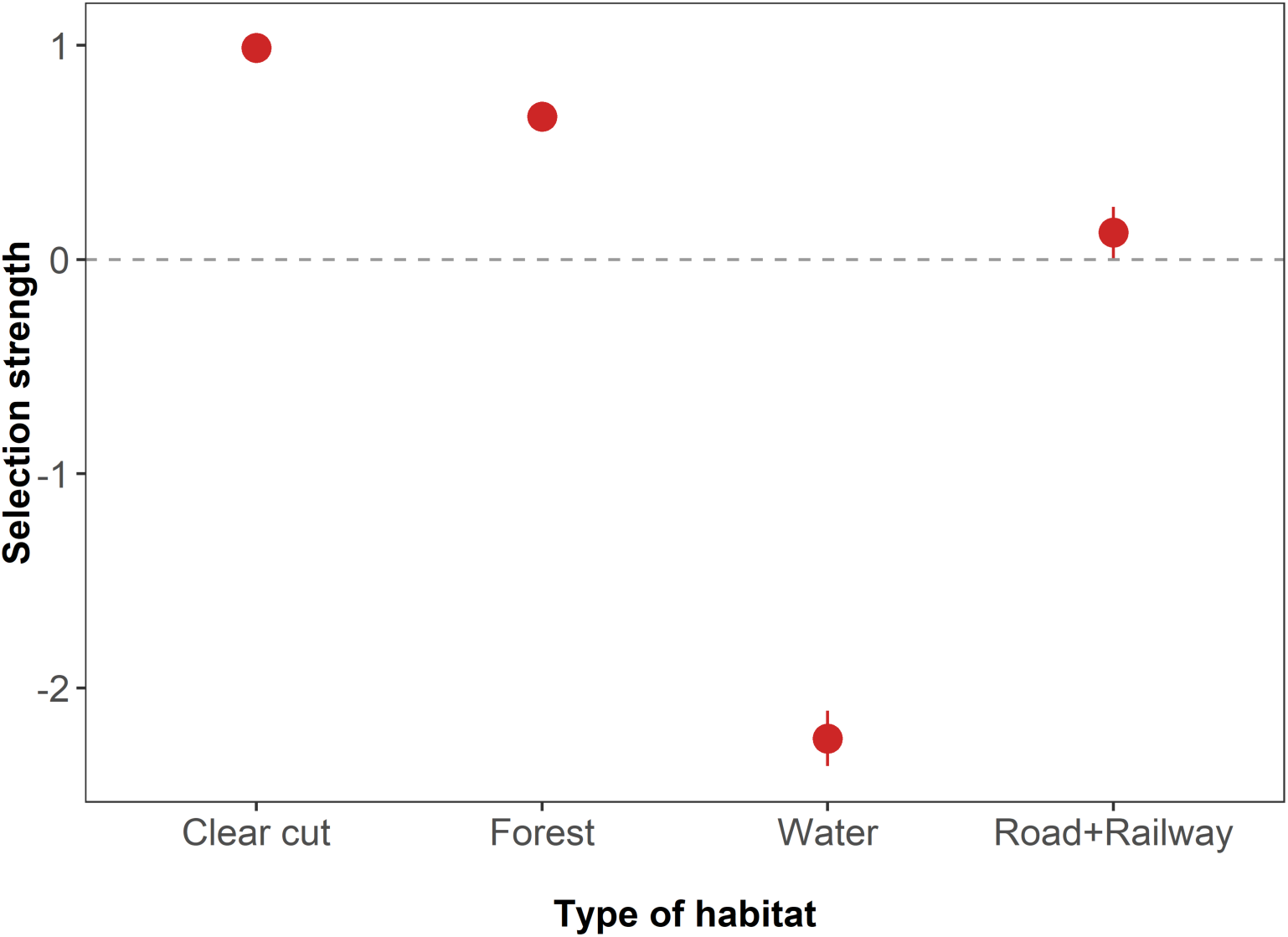
Estimates of integrated step selection function (iSSF) (coefficient ± SE), showing selection of habitat categories by 74 Golden Eagles (both adults and immatures together) in Sweden. Selection of different habitat categories are shown relative to open land (as reference, not shown in the figure). Positive values indicate selection and negative values avoidance.

**Table 1.** Coefficients of the Integrated Step Selection Functions (iSSF) (1 – 14) representing the habitat selection of Golden eagles in relation to linear infrastructure in Sweden using GPS tracking data from 74 individuals (36 Adults and 38 Immatures, n=437217 locations). Positive model coefficients of categorical explanatory variables indicate selection, while for continuous explanatory variables, closeness is indicated by a negative coefficient (models 2 - 14). D_Road_ = distance to nearest road, D_Railway_ = distance to nearest railway, D_Powerline_ = distance to nearest powerline. Age variable is compared between Adults and Immatures (>5 years).

| MODEL | PARAMETER/S | COEFFICIENT | STD.ERROR | Z | P-VALUE |
| --- | --- | --- | --- | --- | --- |
| Habitat Selection |  |  |  |  |  |
| 1 | Road+Railway | 0.13 | 0.06 | 2.04 | < 0.05 |
| 1 | Water | -2.24 | 0.07 | -33.82 | < 0.001 |
| 1 | Forest | 0.67 | 0.02 | 34.65 | < 0.001 |
| 1 | Clear cut | 0.99 | 0.02 | 46.85 | < 0.001 |
| Selection of areas closer to Roads and in seasons |  |  |  |  |  |
| 2 | $D_{\text{Road}}$ | -0.58 | 0.02 | -34.17 | < 0.001 |
| 2 | $D_{\text{Road}}$ :Autumn | -0.38 | 0.05 | -8.13 | < 0.001 |
| 2 | $D_{\text{Road}}$ :Spring | -0.38 | 0.04 | -10.76 | < 0.001 |
| 2 | $D_{\text{Road}}$ :Winter | -0.42 | 0.11 | -3.82 | < 0.001 |
| Selection of areas closer to Railway lines and in seasons |  |  |  |  |  |
| 3 | $D_{\text{Railway}}$ | -0.39 | 0.01 | -31.35 | < 0.001 |
| 3 | $D_{\text{Railway}}$ :Autumn | -0.28 | 0.03 | -8.60 | < 0.001 |
| 3 | $D_{\text{Railway}}$ :Spring | -0.29 | 0.03 | -11.35 | < 0.001 |
| 3 | $D_{\text{Railway}}$ :Winter | -1.16 | 0.13 | -8.73 | < 0.001 |

| Selection of areas closer to Powerlines and in seasons |  |  |  |  |  |
| --- | --- | --- | --- | --- | --- |
| 4 | D<sub>Powerline</sub> | -0.53 | 0.02 | -32.31 | < 0.001 |
| 4 | D<sub>Powerline</sub> :Autumn | -0.36 | 0.05 | -8.13 | < 0.001 |
| 4 | D<sub>Powerline</sub> :Spring | -0.25 | 0.03 | -7.88 | < 0.001 |
| 4 | D<sub>Powerline</sub> :Winter | -0.53 | 0.12 | -4.56 | < 0.001 |
| Search behaviour closer to Roads |  |  |  |  |  |
| 5 | D<sub>Road</sub> | -0.27 | 0.01 | -23.38 | < 0.001 |
| 5 | Log (sl) | 0.11 | 0.00 | 30.29 | < 0.001 |
| 5 | D<sub>Road</sub> :Searching | -0.50 | 0.08 | -6.43 | < 0.001 |
| Search behaviour closer to Railway lines |  |  |  |  |  |
| 6 | D<sub>Railway</sub> | -0.43 | 0.01 | -32.80 | < 0.001 |
| 6 | Log (sl) | 0.09 | 0.00 | 24.96 | < 0.001 |
| 6 | D<sub>Railway</sub> :Searching | -0.25 | 0.06 | -4.07 | < 0.001 |
| Search behaviour closer to Powerlines |  |  |  |  |  |
| 7 | D<sub>Powerline</sub> | -0.14 | 0.01 | -13.29 | < 0.001 |
| 7 | Log (sl) | 0.11 | 0.00 | 30.53 | < 0.001 |
| 7 | D<sub>Powerline</sub> :Searching | -0.05 | 0.04 | 1.22 | 0.22 |
| Sitting behaviour closer to Roads |  |  |  |  |  |
| 8 | D<sub>Road</sub> | -0.89 | 0.04 | -24.96 | < 0.001 |
| 8 | Log (sl) | 0.03 | 0.00 | 14.08 | < 0.001 |
| 8 | D<sub>Road</sub> :Sitting | -1.13 | 0.05 | -23.16 | < 0.001 |
| Sitting behaviour closer to Railway lines |  |  |  |  |  |
| 9 | D<sub>Railway</sub> | -0.98 | 0.04 | -36.51 | < 0.001 |
| 9 | Log (sl) | 0.02 | 0.00 | 8.06 | < 0.001 |
| 9 | D<sub>Railway</sub> :Sitting | -0.84 | 0.04 | -20.93 | < 0.001 |
| Sitting behaviour closer to Powerlines |  |  |  |  |  |
| 10 | D<sub>Powerline</sub> | -0.48 | 0.04 | -13.41 | < 0.001 |
| 10 | Log (sl) | 0.03 | 0.00 | 13.71 | < 0.001 |
| 10 | D<sub>Powerline</sub> :Sitting | -0.95 | 0.05 | -19.57 | < 0.001 |
| Selection of areas closer to Roads by Age class |  |  |  |  |  |
| 11 | D<sub>Road</sub> | -0.53 | 0.02 | -29.88 | < 0.001 |
| 11 | D<sub>Road</sub> :AgeImmature | -0.41 | 0.03 | -15.15 | < 0.001 |
| Selection of areas closer to Railway lines by Age class |  |  |  |  |  |
| **12** | D<sub>Railway</sub> | -0.29 | 0.01 | -22.00 | < 0.001 |
| **12** | D<sub>Railway</sub> :Age Immature | -0.42 | 0.02 | -21.68 | < 0.001 |
| Selection of areas closer to Roads by Age class and number of years |  |  |  |  |  |
| **13** | D<sub>Road</sub> Immature | -0.85 | 0.04 | -20.57 | < 0.001 |
| **13** | D<sub>Road</sub> Immature:years | -0.05 | 0.02 | -3.12 | < 0.01 |
| Selection of areas closer to Railway lines by Age class and number of years |  |  |  |  |  |
| **14** | D<sub>Railway</sub> Immature | -0.44 | 0.03 | -14.44 | < 0.001 |
| **14** | D<sub>Railway</sub><br>Immature:years | -0.16 | 0.01 | -12.17 | < 0.001 |

There was a consistent spatial trend that eagle traffic accidents were observed throughout Sweden (Figure 1a & 1c>). This was also ascertained from the study eagles, where six individuals were killed by traffic and five are visible in the map (Figure 3). Figure 4 shows examples of the range of trap habitat encountered at a landscape scale including locations and type of position during annual movement paths of birds. It shows that eagles encounter linear features multiple times across their annual movements and face the risk of collisions throughout their range.

**Figure 3.**
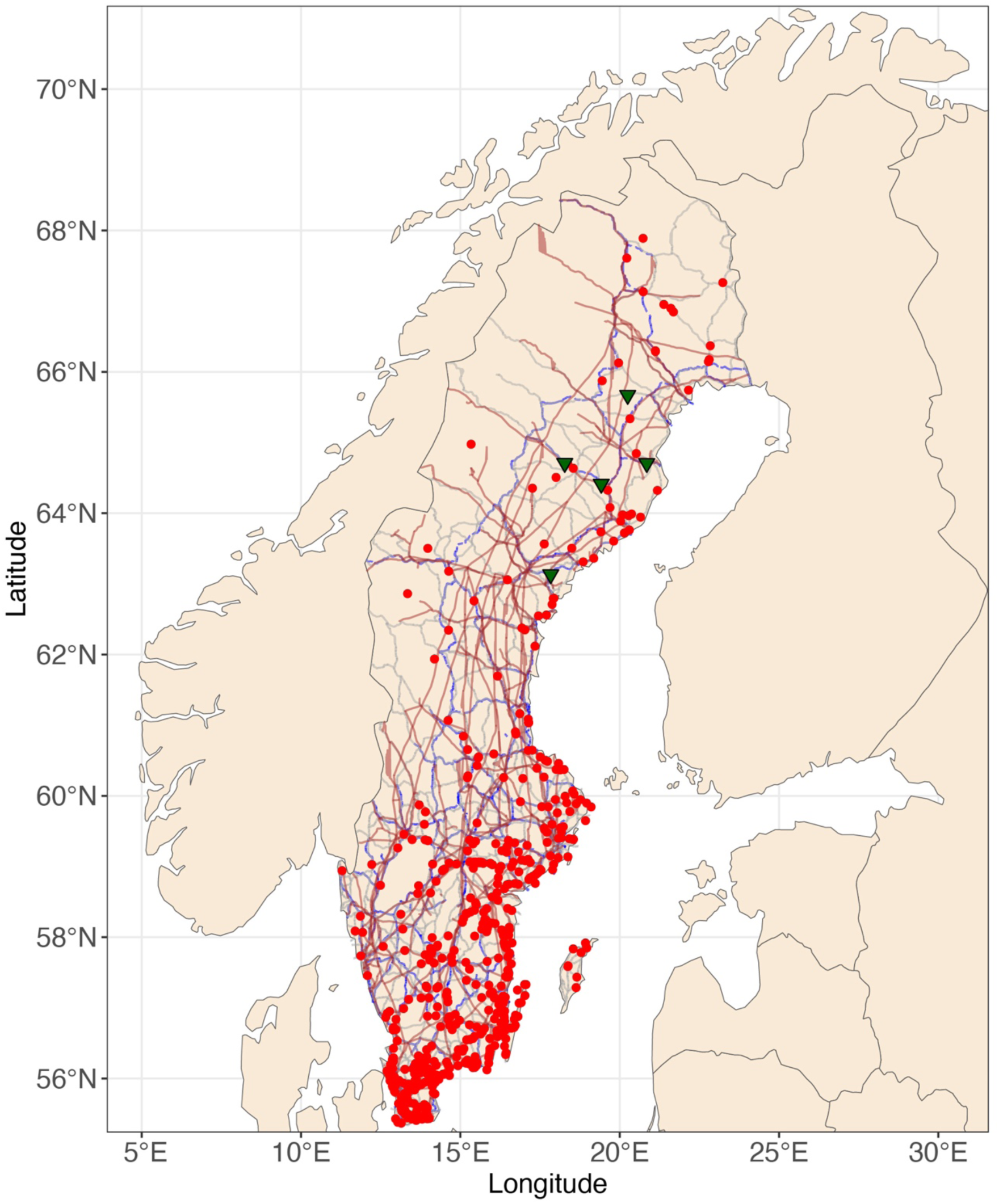
Locations of Eagle Traffic collisions (Red dots) recorded throughout Sweden during 2010-2023 (n=650) by the SNRAD. Green triangles represent the locations of collisions of five GPS marked Golden eagles with traffic. Only Primary Roads are shown, represented in grey, Railway lines in blue and the main Powerlines in brown.

**Figure 4a, b & c.**
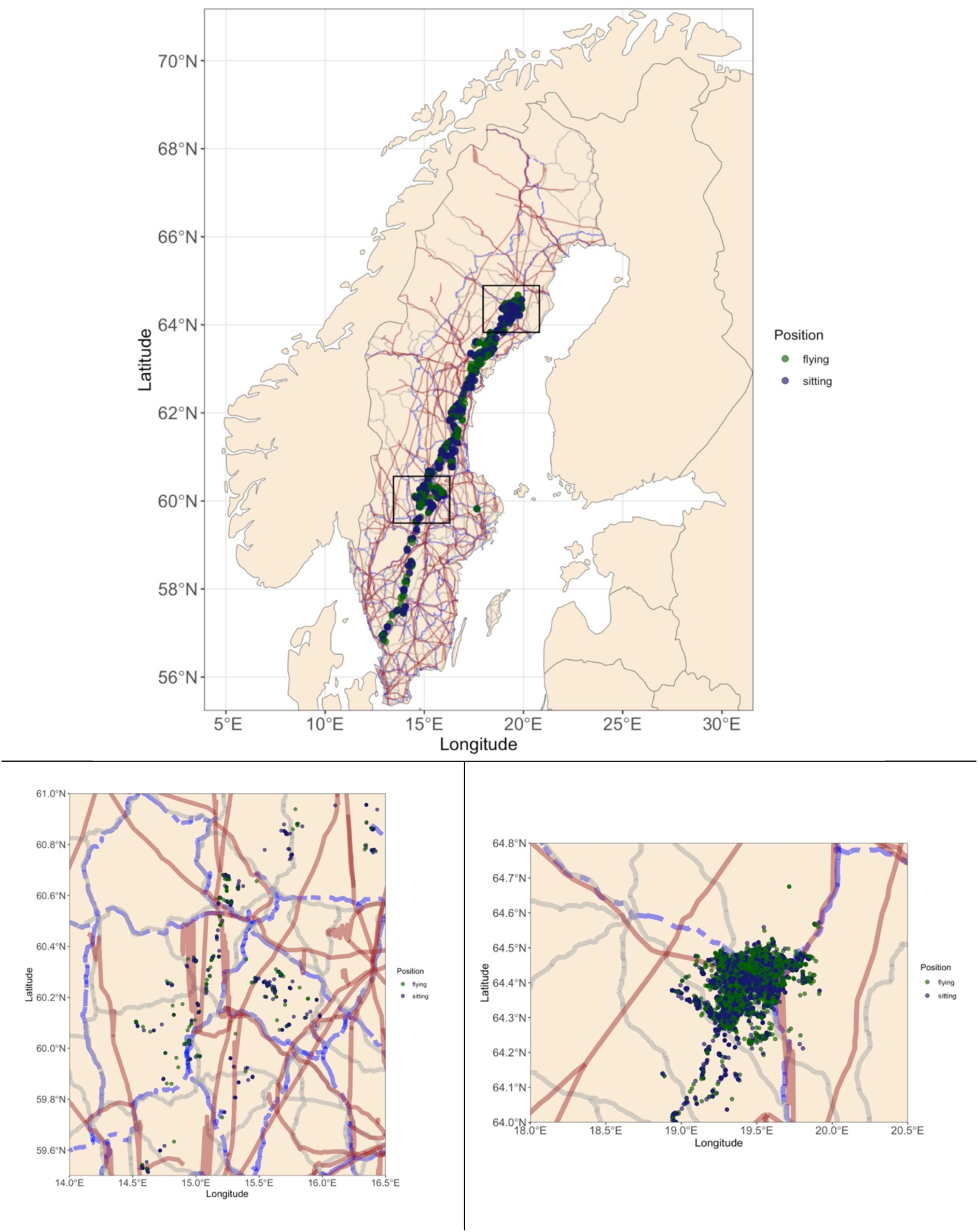
a) Example annual movement track of an Immature Golden eagle in Sweden with labelled sitting and flying positions to demonstrate interactions with linear infrastructure. b) Magnified view of part of the track of the same individual when in southern Sweden, c) and same individual when in northern Sweden. Only Primary Roads are shown, represented in grey, Railway lines in blue and the main Powerlines in brown.

**Figure 4d, e & f.**
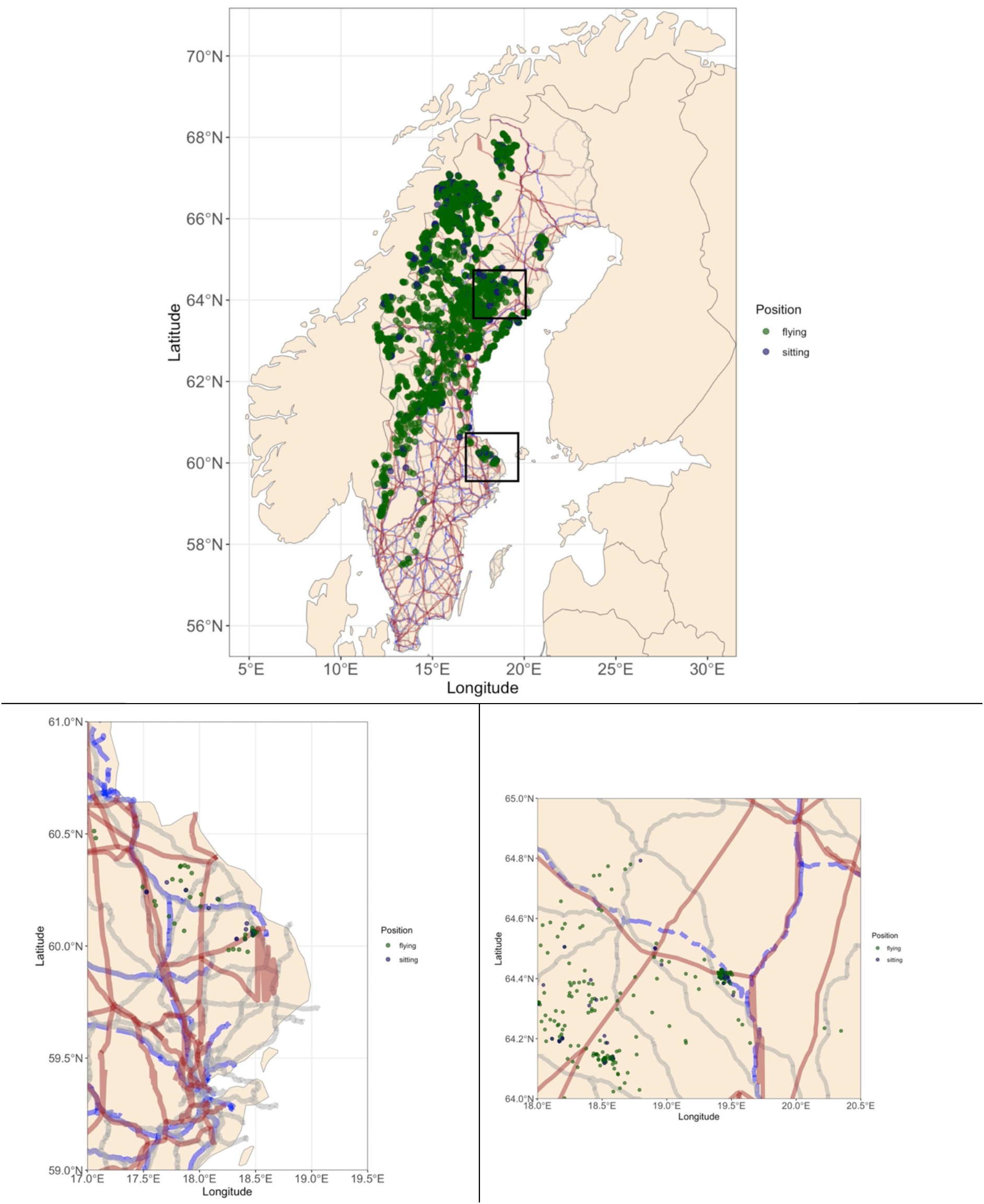
d) Example multiannual movement track of an Immature Golden eagle in Sweden with labelled sitting and flying positions to demonstrate interactions with linear infrastructure. e) Magnified view of part of the track of the same individual when in southern Sweden, f) and same individual when in northern Sweden. Only Primary Roads are shown, represented in grey, Railway lines in blue and the main Powerlines in brown.

**Figure 4g, h & i.**
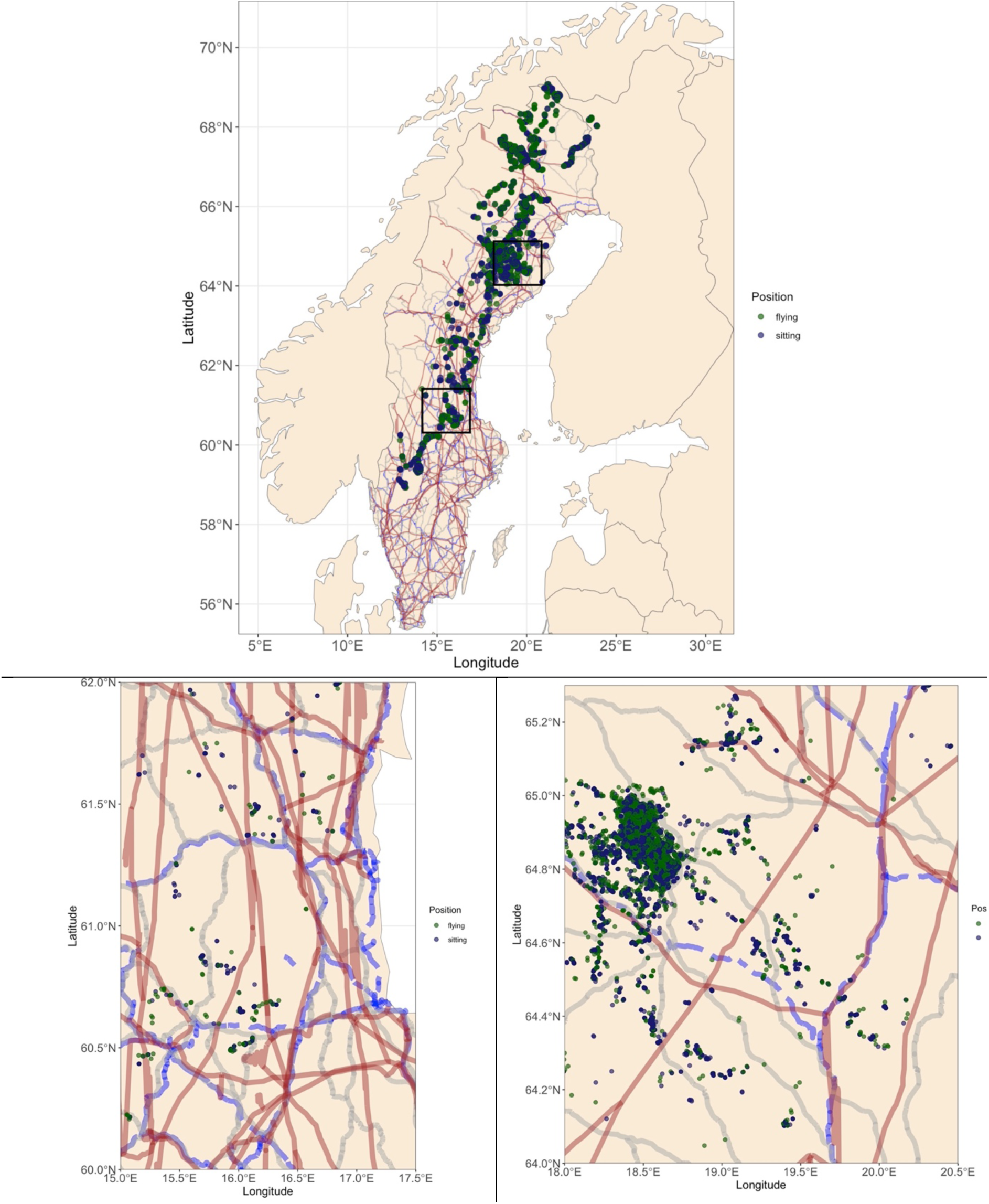
g) Example multiannual movement track of an Immature Golden eagle in Sweden with labelled sitting and flying positions to demonstrate interactions with linear infrastructure. h) Magnified view of part of the track of the same individual when in southern Sweden, i) and same individual when in northern Sweden. Only Primary Roads are shown, represented in grey, Railway lines in blue and the main Powerlines in brown.

The temporal seasonal trend of wildlife accident reveals an increasing number of accidents since 2010 (Figure 5a & b, Figure S3) and annually towards winter, with two general peaks (Figure S2- Appendix). One peak occurs in May and June and the other occurs during September to December. Overall, the trend provides a mean estimate of number of accidents per month as an index of attractiveness of roads and railways.

### Behaviour in trap habitat

Close to roads and railways, individuals performed more search than travel behaviour, while at powerlines there was no indication for search movements (Table 1, β_road_ = - 0.50 ± 0.08, β_railway_ = - 0.25 ± 0.06, β_powerline_ = - 0.05 ± 0.04). Additionally, eagles were sitting more frequently instead of flying at linear infrastructure (Table 1, β_road_ = - 1.13 ± 0.05, β_railway_ = - 0.84 ± 0.04, β_powerline_ = - 0.95 ± 0.05).

### Selection of trap habitat and learning by immatures

Immature eagles were significantly closer to roads and railways compared to adults (Table 1, β_road_ = - 0.41 ± 0.03, β_railway_ = - 0.42 ± 0.02). As immature eagles got older, they moved closer to roads and railways (Table 1, β_road_ = - 0.05 ± 0.02, β_railway_ = - 0.16 ± 0.01).

## Discussion

In this study, through an extensive and unique multi-annual dataset, we demonstrate strong indication that linear infrastructure creates an ecological trap (ET) for Golden Eagles across space and time. Among the conditions that need to be fulfilled to demonstrate an ecological trap, eagles consistently selected, searched and sat frequently closer to roads and railways, called ‘trap habitats’ here, across seasons, years and throughout their range in Sweden. The negative demographic consequences at the population level, could be inferred through our data and published studies, e.g. Ecke et al. 2017, that show the accidents and trauma being the dominant known source of mortality. The SNRAD includes the observed instances of dead eagles, thus ascertaining the demographic costs of foraging in these habitats. The increasing mortality numbers over years (Figure S4) also reflect the effort and improvement in monitoring system and awareness over time besides the increase in accidents.

Eagles in the boreal landscape are known to depend on cyclic prey species like mountain hare (*Lepus timidus*), grouse species (*Lagopus spp.*) and many species of medium sized and small mammals (Moss, 2015; Watson, 2010). These species are forest dwelling and are represented in the simultaneous selection of coniferous forests and clear cuts in our analyses (Singh et al., 2017). However, many of these species are currently suffering the negative impact of climate change, over harvesting and human modifications of landscapes, on their population demography and many are believed to have lost their cyclicity under the influence of these human driven changes in ecosystems (Cornulier et al., 2013). This likely has cascading impacts on eagle prey selection, behaviour and demography through their increased dependence on carrion, visible in the selection of attractive trap habitats. We observed consistent selection for trap habitats across the landscape, seasons and years, which implies that this is already occurring and eagles may have learned to compensate for the lower abundances of prey species by switching to carrion. Over a short-term (5-10 years) there might be a positive impact on eagle population through increased recruitment and reproductive output linked to higher food availability and turn over, but over a longer term (> 10 years) this may have a catastrophic impact considering a high mortality of prime aged adults and immatures from traffic accidents.

We observed that individuals selected and were observed sitting more frequently along powerline corridors, but in contrast to roads and railways, eagles did not show search behaviour around powerlines. This suggests that powerline poles are likely functioning as perching sites to hunt and scan for scavenging opportunities from remains of other electrocuted or collision- killed birds in powerline corridors. It explains why the eagles face electrocution risk as reported in other studies (Janss, 2000; Krüger et al., 2004; Slater & Smith, 2010). For e.g. Golden eagle mortality due to electrocution is at very high levels in the United States (Ansell & Smith, 1980; Harness & Wilson, 2001). This could be due to the kind of poles and material used for powerline poles and distance between the tops and the high tensions wires that facilitate a higher contact between the wires and the sitting birds. Alternatively, a higher food availability within the powerline corridors and resulting frequent use by the eagle drives these patterns. Some data on number of electrocutions has been earlier presented by Ecke et al. (2017).

The extent and amount of wildlife traffic accidents in Sweden have been steadily increasing and are currently at all-time high. Within the years 2015 to 2023, the wildlife accidents have increased from c. 45000 individual reported accidents to c. 69000 (www.viltolycka.se), and these include only the larger species (e.g. deer species, brown bear, lynx, wild boar and eagles) which are required to be reported by law. It is possible that many other scavenger species such as wolverine (*Gulo gulo*), ravens (*Corvus corax*), red fox (*Vulpes vulpes*), brown bear (*Ursus arctos*) are facing a similar ecological trap situation as eagles, from roads and railroads because of the increase in accidents of other species. In fact, other human activities such as garbage creation, livestock rearing and other feed subsidies are creating similar situations elsewhere (Oro et al., 2013). Grizzly bears (*Ursus arctos*) in urban environments have been reported to be in an ecological trap due to their dependence on human created garbage dumps (Lamb et al., 2017). The vulture crisis in Asia caused by poisoning from Dichlofenac through livestock carcasses driving > 95% decline in populations is a prominent example of an ecological trap (Prakash et al. 2003). Pumas (*Puma concolor*) have been shown to kill livestock prey available through humans, and in retaliation face risk of being killed, due to a lower perception of risk from humans (Nisi et al., 2022). We also know that the other eagle species in the region, the White-tailed Sea eagle (*Haliaeetus albicilla*), which has an even greater tendency to scavenge, is facing a similar situation, with 60% of the mortality attributed to human related factors (Isomursu et al., 2018). Such diversity of examples indicates that even the same species might encounter multiple trap situations throughout its annual life cycle.

Eagles in our study system are already exposed to a number of other human induced rapid environmental changes besides wildlife traffic accidents that may create multiple ecological trap situations. These include ongoing rapid wind farm development, conflicts with reindeer and sheep husbandry through predation and retaliatory protective hunting, and lead poisoning from consumption of ammunition infested carcasses left over in the forest from annual moose hunt (Ecke et al., 2017; Singh et al., 2021). Indeed, these all have been reported as main causes for eagle mortality in Sweden and even Finland, besides the traffic collisions (Ecke et al. 2017). In our study, the only individuals where the cause of death could be confirmed was through the traffic accidents and when the last known position was along a road or railroad. In some other cases when the fate of the bird was unknown, we often attributed that to transmitter failure, as many birds were seen to be alive even after the transmitter stopped functioning. Another study showed the relationship between lead poisoning and consequent changes in eagle movement behaviour and flight capacity, leading to a higher number of collisions and increasing the likelihood of death by more than three folds (Ecke et al. 2017, Isomursu et al. 2018). This shows many indirect interactions between human induced environmental changes and their cumulative impact on species populations potentially leading to a severe trap situation from combined mortality. All of this presents a rather grim picture for eagle population in the region.

Literature suggests that due to a lack of natural predators, apex consumers lack the capacity to perceive novel sources of risk, especially human induced rapid land use modifications, which increases their vulnerability to ecological traps (Ripple et al., 2014). Fast moving traffic and windmills are such novel risks and unless carefully monitored and reported, the actual levels of mortality are very difficult to quantify in these sites and for wide ranging species such as eagles. What we have in this study is probably a very small proportion of the mortality found and reported and we have no estimate of age of the dead birds. Attractive food subsidies through traffic accidents may keep the immature individuals within traps and prevent dispersal, increase mortality, reduce their ability to hunt or compete for mates, food and territories (Lamb et al. 2017). A high female mortality may also mean that many females may be too young to reproduce leading to low recruitment in the population and low dispersal out of the trap (Proctor et al., 2012). Future monitoring programmes at wind farms and infrastructure sites should therefore incorporate careful monitoring of demographic parameters such as age and species of birds. Similarly, other interacting factors such as lead ammunition use in hunting should be banned in parallel. Indeed, the lead levels in liver of traffic killed eagles in Sweden has been shown to be significantly high (Ecke et al. 2017).

Considering the fact that distribution of traffic accidents is widespread, hunting with lead ammunition is also at the same scale, wind farm development is occurring throughout the country both on land and offshore, all individuals are similarly exposed to these threats because of their migratory movements across a north-south latitudinal gradient (Singh et al. 2017). This presents a challenge which requires careful national and regional coordination of mitigation measures such as carcass removal after accidents, careful monitoring and reporting of collisions, innovations in warning systems for both the wildlife and humans, among others. It also requires coordination between actors and stakeholders such as the traffic authorities, environmental protection agency, hunting community, industry and other environmental policymakers. It is clear that there are certain wildlife traffic collision hotspots for eagles in Sweden which can be focussed for mitigation measures right away (Figure S5 - Appendix). This is also true for other species that are being involved. Combining data from accidents and eagle space use via GPS can be the foundation to build mitigation measures.

The eagle populations have only recently recovered after the Europe wide ban in DDT (Hailer et al., 2006; Korsman et al., 2012) but are now being exposed to high levels of novel environmental contaminants (Dürig et al., 2022; Sturm & Ahrens, 2010). To conclude, there is an urgent need to eliminate the discussed threats while handling emerging ones simultaneously and enhancing population monitoring of key demographic parameters. We suggest that future studies should therefore incorporate multiple potential traps and other threats simultaneously to understand the impact of anthropogenic changes on species populations and ecosystems to ensure and improve conservation.

## Acknowledgements

The project was financed by the Vindval programme of the Swedish Environmental Protection Agency, 2010. NS was financed by the EU Horizon Programme 2020 - SUPERB (Systemic solutions for upscaling of urgent ecosystem restoration for forest-related biodiversity and ecosystem services) Grant agreement ID: 101036849.

The Authors have no competing interests.

## Author Contributions

NS & ME Conceptualized the MS. NS, FE and BH organized Funding and Data curation. ME and NS conducted the analyses. NS and ME wrote the first draft of the MS and all authors contributed to writing and interpretation of data.

## Appendix

**Figure S1.**
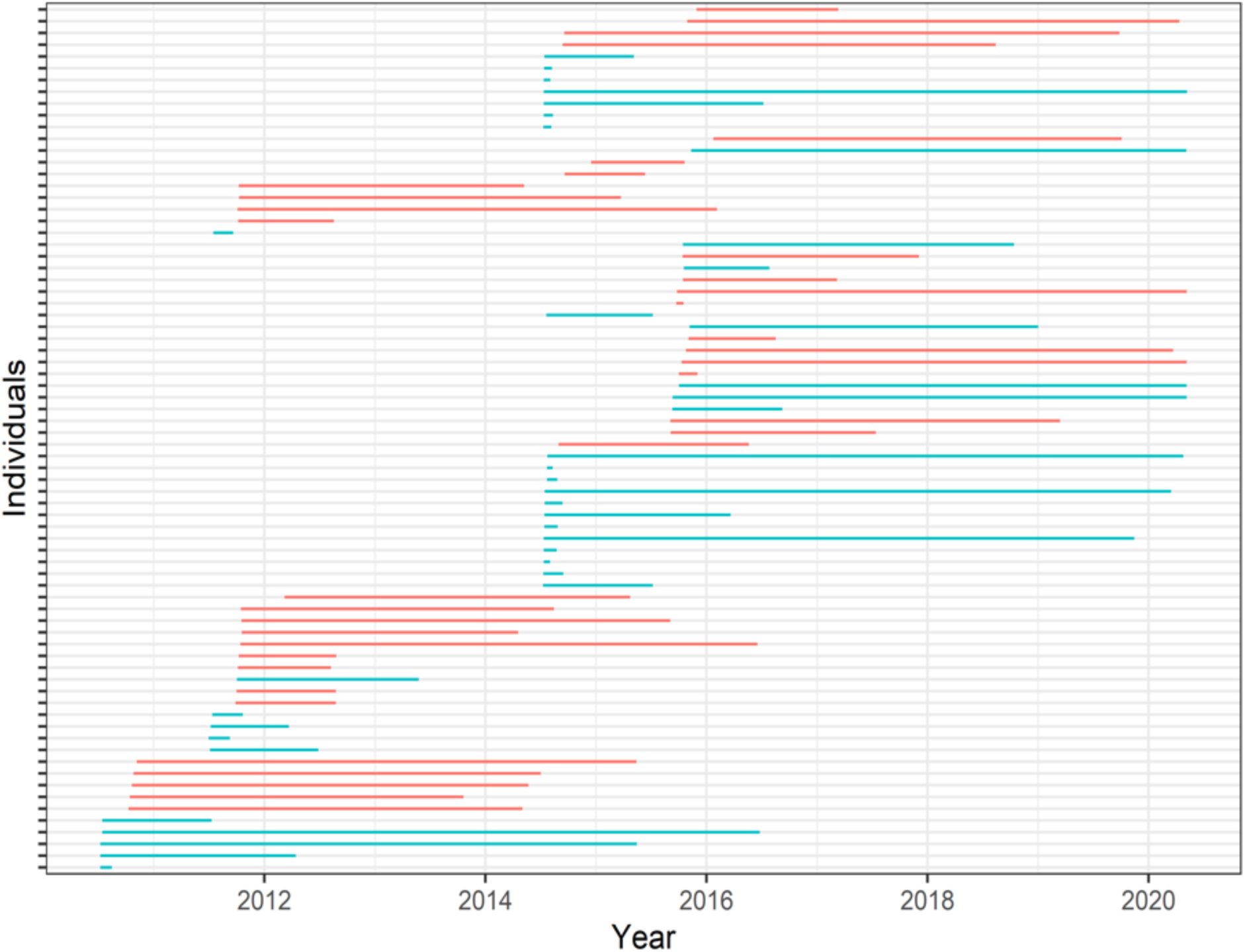
Individual tracking periods of 36 Adult Golden eagles (in red) and 38 Immatures (in blue). Each line indicates one individual.

**Figure S2.**
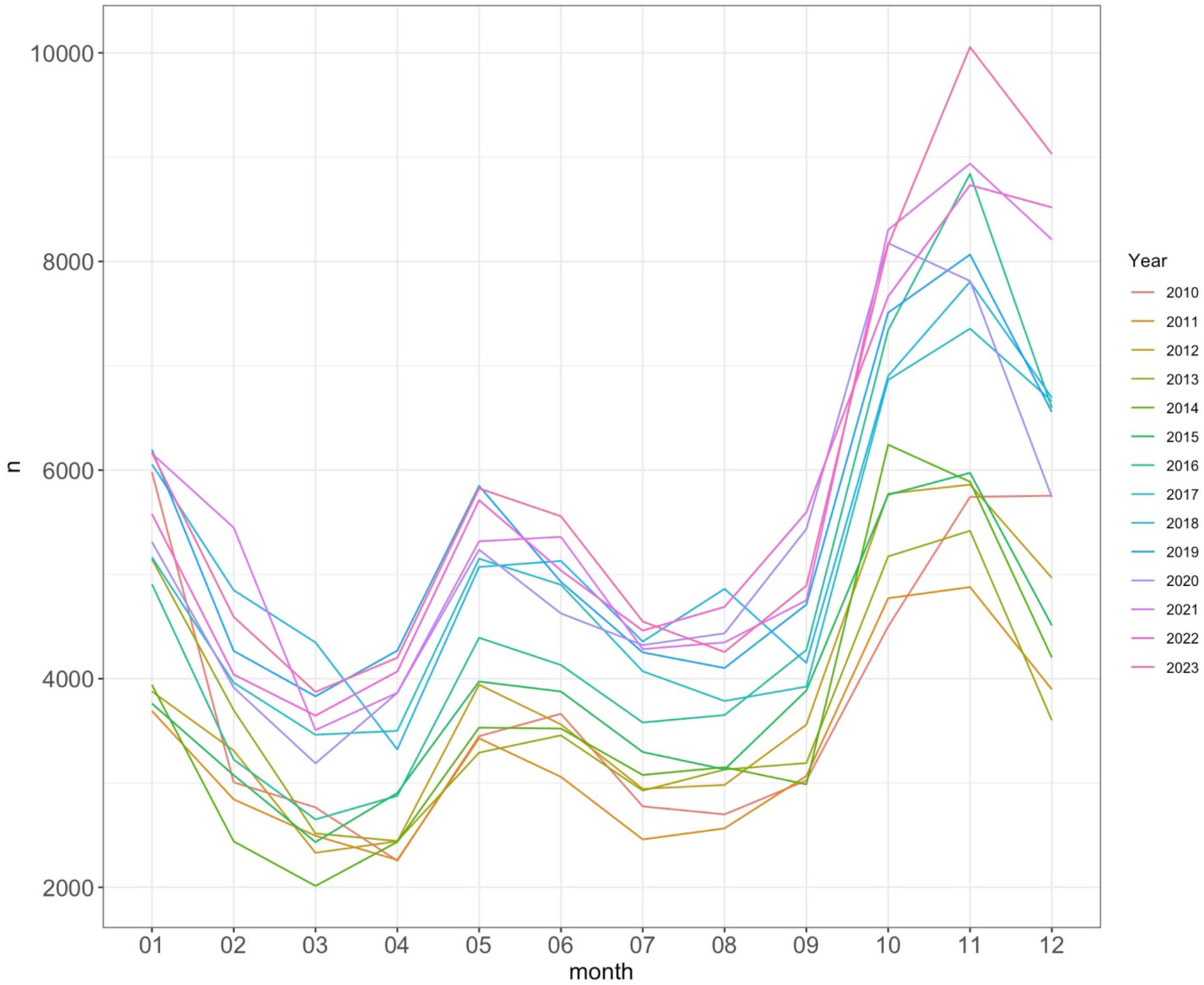
Mean number of wildlife traffic accidents of selected species (Moose, Roe deer, Red deer, Fallow deer, Wild boar, Bear, Wolf, Lynx, Wolverine and Otter together) in Sweden by month during the study period 2010-2023. SD was not included in the figure due to high variation in mean number of accidents among species. For more details see Methods. Data obtained from SNRAD.

**Figure S3.**
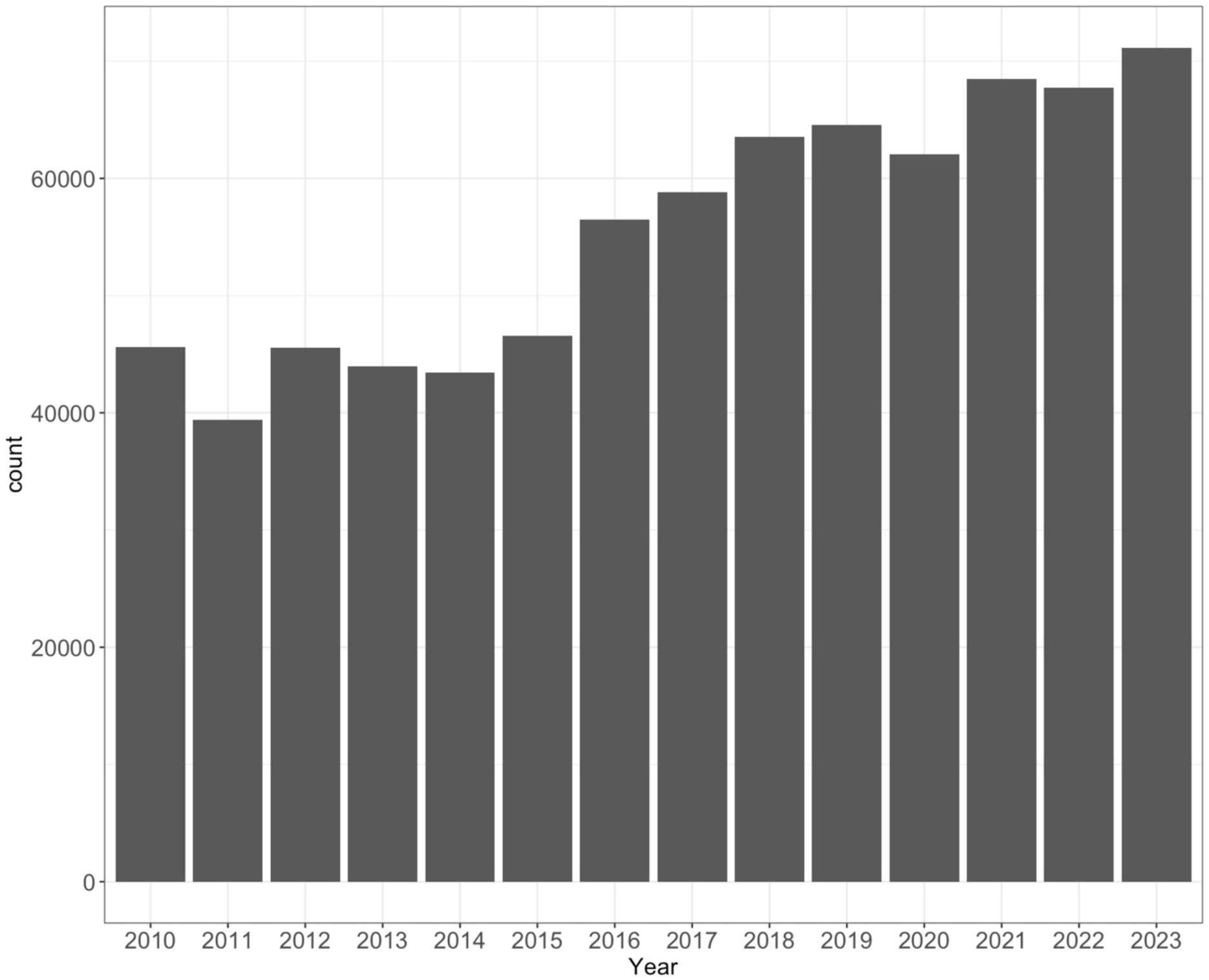
Total number of reported wildlife traffic accidents (for Moose, Roe deer, Red deer, Fallow deer, Wild boar, Bear, Wolf, Lynx, Wolverine and Otter) in Sweden during 2010 – 2023. For more details see Methods. Data obtained from the SNRAD. www.viltolycka.se

**Figure S4.**
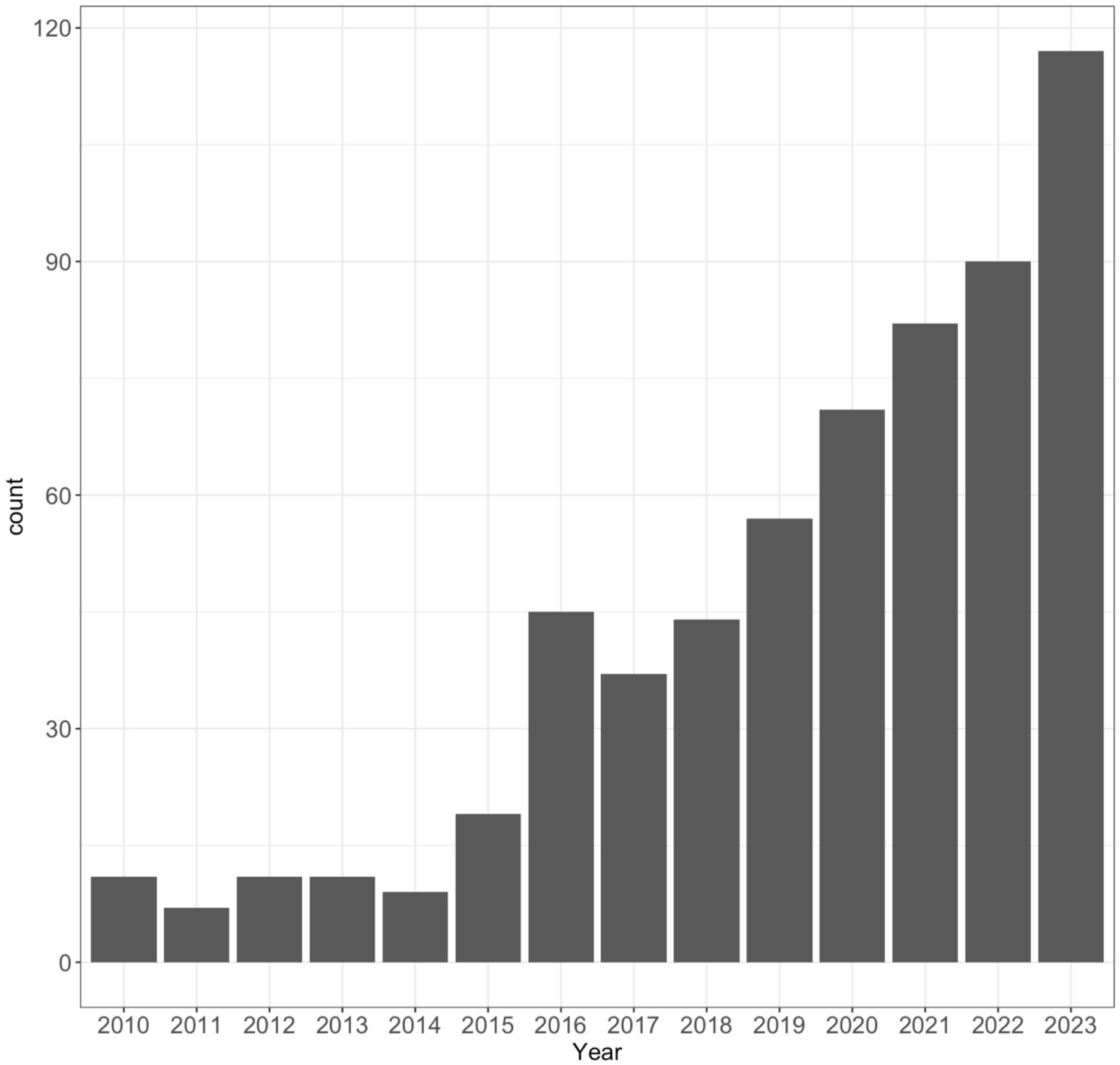
Total number of Eagle traffic accidents in Sweden during 2010 – 2023. For more details see Methods. Data obtained from the SNRAD. www.viltolycka.se

**Figure S5.**
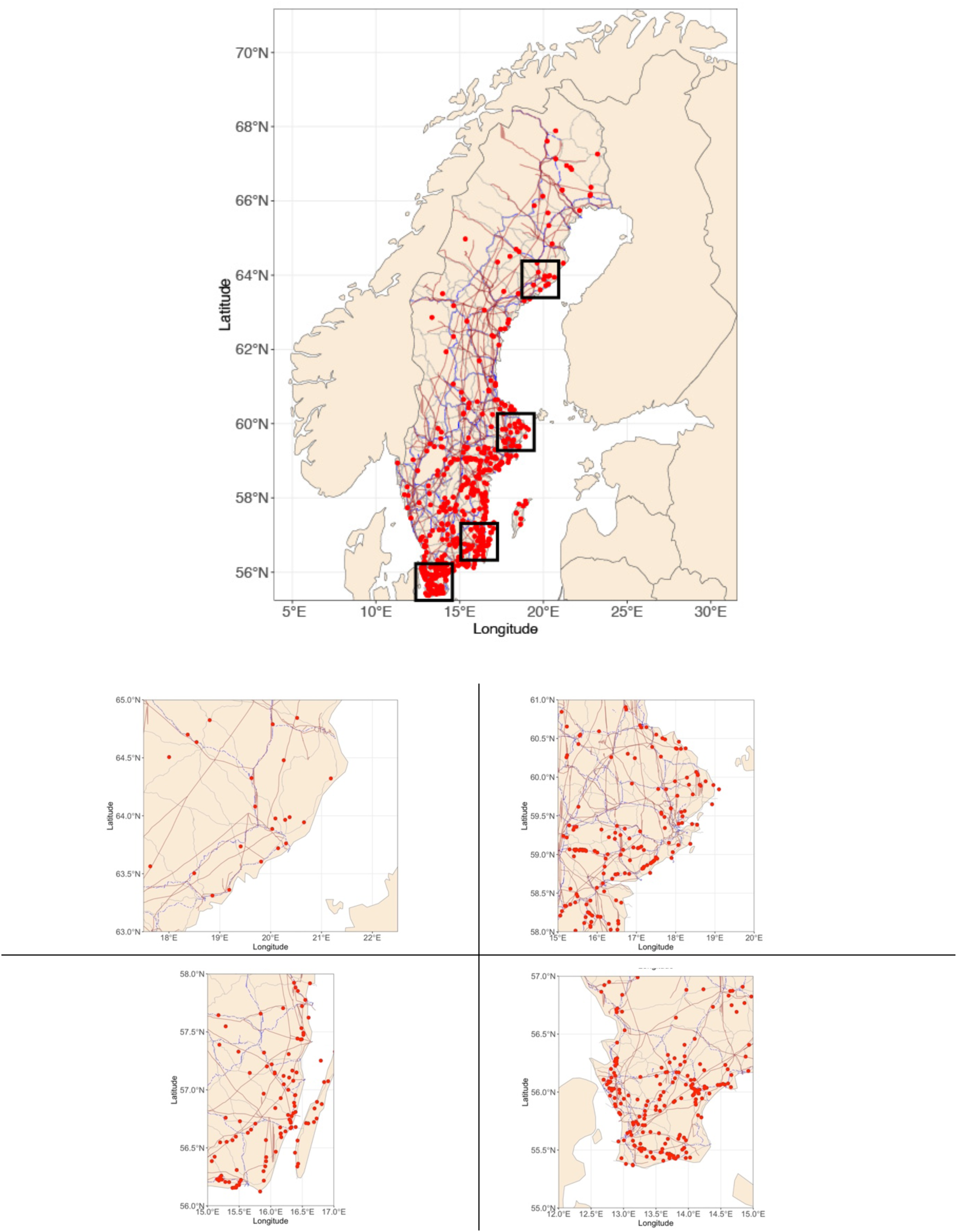
Zoomed out views of four eagle traffic accidents hotspots across Sweden which could be focussed for immediate mitigation and management plans by the authorities.

